## Supplementary Material for "Speos: An ensemble graph representation learning framework to predict core genes for complex diseases"

### Supplementary Notes

#### Supplementary Note 1: Biases and PPI network selection

To estimate how the data generation process of different networks may affect predictions, we compared the topological measures for positively labeled Mendelian and the remaining genes. In systematically acquired large-scale datasets networks such as HuRI[18] and BioPlex[19], Mendelian disorder genes have almost the same probability to be in the neighborhood of other Mendelian disorder genes as unlabeled genes (**Extended Data Fig. 1a**). In contrast, in networks generated by agglomerating data from hypothesis-driven approaches the proportion of other Mendelian disorder genes in their neighborhood (homophily) increases. This drastic discrepancy reflects on one hand the well documented inspection bias of hypothesis-driven approaches[18,25,26], but may also be influenced by the incompleteness of high-throughput approaches (**Extended Data Fig. 1b**). The degree distribution of Mendelian and unlabeled genes further confirms the substantial inspection bias in the small-scale literature. While there is no meaningful difference in the distribution and mean degree of both groups of genes in the tested unbiased networks, in networks based on the small-scale literature, Mendelian disorder genes show a substantial shift towards higher degrees than unlabeled genes, this reflecting the ‘interest’ of the community in specifically studying these genes (**Extended Data Fig. 1c**). Since GNNs are influenced by the label distribution in the direct neighborhood[118], this effect of publication bias on the neighborhood label distribution might artificially increase performance on a holdout set while at the same time perpetuating existing biases if hypothesis-driven networks are used. Although IntAct[22] shows biased patterns towards a higher mean degree of Mendelian disorder genes, we include it in our analyses both as a reference to other studies and to account for interactions that are not covered by high-throughput approaches, but we monitor the effect of its biases on the resulting candidate genes (**Supplementary Note 4**). STRING[23] (cutoff 0.7), on the other hand, is excluded due to its drastic difference in degree distributions between Mendelian disorder genes and unlabeled genes.

**Supplementary Note 2: Design and optimization of GNN**

Our model consists of three parts, pre-message passing (pre-MP), message passing (MP) and post-message passing (post-MP) (**Extended Data Fig. 4**). Pre- and post-MP are MLPs built from stacked fully connected layers interspersed with ELU nonlinearity, while the MP module is built from GNN layers, interspersed with ELU nonlinearity and instance normalization layers. The pre-MP module is built of one input layer which transforms the input space into the hidden dimension $dim_{hid}$ of our model and $q$ further layers which extract node-level features and feed it into the MP module. We use the same hidden dimension $dim_{hid}$ across all hidden layers of the model. The MP module is built from $r$ blocks each consisting of one GNN, one nonlinearity, and one normalization layer, which then feeds the latent features into the post-MP module. The post-MP module consists of $s$ fully connected layers for node-level pattern recognition and two fully connected layers to transform it into the one-dimensional output space with an intermediate step of $\frac{dim_{hid}}{2}$ dimensions. We have searched hyperparameters $q$ and $s$ for all iterations from 1 to 6 without much difference in performance (data not shown). Therefore we settled at $q = s = 2$, which leaves both modules ample room for feature recognition and keeps the number of model parameters small. When evaluating the impact of the dimensionality of the pre- and post-MP input/output vectors $dim_{hid}$ at 10, 20, 30, 40, 50, 75, 100, 125 we observed no performance change exceeding one standard deviation of the AUROC; we fixed $dim_{hid}$ = 50. Although hidden dimensions larger than 125 were included in the code, the runs went out of memory on a 40GB A100 GPU for the large, multi-network models and did not return results.

For the MP module, we tested 35 adjacency matrices and eleven GNN layers from the repertoire of PyTorch Geometric, of which eight are single-network GNNs, which only use a single adjacency matrix and which are evaluated first, and three are multi-network GNNs, that can use multiple adjacency matrices simultaneously. Setting the number of GNN layers to $r = 0$ collapses the model into an MLP with only pre-MP and post-MP and renders the influence of adjacency matrix and GNN layer to zero. **Extended Data Fig. 5** shows the influence of different single-network GNN layers and the hyperparameter $r$ on the performance of the model using the adjacency matrices BioPlex 3.0 HEK293T, GRNdb Adipose Tissue and IntAct Direct Interaction. We evaluated the topology adaptive graph layer (TAG)[47], GraphSAGE[131], the graph transformer layer[132], Chebyshev graph convolution layer[133], the seminal graph convolution layer (GCN)[46], simplified graph convolution layer (sGCN)[134], the graph attention layer (GAT)[135] and the graph isomorphism layer (GIN)[136]. We conducted the hyperparameter optimization for gene labels of immune dysregulation (**Extended Data Fig. 5**) and cardiovascular disease (now shown). For two adjacency matrices, most layers reduce the model performance compared to an MLP ($r = 0$). Only with IntAct Direct Interaction (**Extended Data Fig. 5c**), the GNN ($r > 0$) outperforms the MLP. However, across all three adjacency matrices, the TAG layer usually outperforms all other layers. The best performing layers, especially according to **Extended Data Fig. 5b, c** are TAG, GraphSAGE and the graph transformer layer. These layers either have mechanisms to block out the neighborhood and retain the self node's features during the convolution (TAG and GraphSAGE) or have a non-linear neighborhood attention mechanism (transformer), allowing them to modulate the incoming message. Simpler GNN layers, however, fail to perform compared to the MLP, which hints towards the tendency of the full neighborhood information being harmful for prediction. From these data we settled for 2 TAG layers which generally achieve the best performance and do not deteriorate performance below the performance of an MLP. After fixing the MP module to 2 layers of TAG, we investigated the performance on all 35 adjacency matrices. In this analysis we also included three multi-network GNN layers that use all adjacency matrices which are typed and fed in the network simultaneously: RGCN[48], RGAT[113] and FiLM[49]. RGAT went out of memory for all runs on a 40GB A100 GPU and is not shown. As recent analyses[137] showed that allowing the information flow to also bypass the GNN can drastically improve the performance, we also evaluated a skip-connection from pre-MP to the output of MP and concatenation of pre-MP's and MP's output features prior to post-MP (**Extended Data Fig. 4c**). From the single-adjacency runs, only Intact Direct Interaction and Intact Physical Association provide a benefit, likely reflecting the above discussed bias (**Extended Data Fig. 6a)**. Using all adjacency matrices simultaneously is only helpful if the FiLM layer is used. Furthermore, using a skip-connection or concatenation is not beneficial, possibly because the TAG layer already contains skip-connections and the FiLM layer is flexible enough to block and override unhelpful information. We therefore settle on using the FiLM layer with all adjacency matrices and the TAG layer with IntAct Direct Interaction without any further skip-connections or concatenations.

Based on the HPO we have settled on two combinations of GNN layer and network: 1. using $r = 2$ TAG layers on the IntAct Direct Interaction network and 2. using $r = 2$ FiLM layers on a merge of all networks. Due to the strict annotation criteria the IntAct Direct Interaction network is very sparse (**Extended Data Fig. 6b**) and most nodes are isolated, whereas after merging all networks for the FiLM layer, only 33 nodes are not in the large connected component (**Extended Data Fig. 6c**). The average shortest path length between positives and between all genes are much shorter in the combined network than in IntAct Direct Interaction. However, in both graphs positives and unlabeled nodes do not differ in their average shortest path lengths. Also, when combining the networks, the node degree distribution shifts away from a scale-free graph as low-degree nodes become less frequent than mid-degree nodes. Finally, positives in Intact Direct Interaction have a much higher frequency of other positives in their neighborhood than when all networks are combined.

**Supplementary Note 3: Network features influencing model performance**

Several observations indicate that, while the input features of every node and adjacent edges are important for its own prediction, the features of neighboring nodes may be irrelevant or even harmful for the machine learning task. In the initial evaluation of base classifiers (**Fig. 1a)**, the best performing method for recovering held out positives is N2V+MLP. This method first uses a random-walk based neural network to project the graph topology into vector space without any node features. These vectors, which describe the position of nodes in the graph, are concatenated with the regular input features like GWAS summary statistics and gene expression for the node of interest and feed it into an MLP. Since the MLP part of N2V+MLP processes every gene in isolation, the neighboring node's features are not seen by the model. It only processes the input features of the node itself along with the topological information of its position in the graph, encoding its connections to other nodes. At the same time we established that this topological information is valuable as N2V+MLP outperforms an MLP that only uses the regular input features in the recovery of held out positives (**Fig. 1a**) and in the discovery of unlabeled positives (**Fig. 3a**). In contrast to N2V+MLP, GNNs process graphs by convoluting the features of adjacent nodes along the connecting edges using the message passing framework. Thus, the edges are used to determine "where" the convolution is applied, but the neighboring nodes' features define the content that is processed and propagated through the network.

We observed that GNNs are outperformed by N2V+MLP in recovering held out positives (**Fig. 1a**), but perform better or comparable to N2V+MLP in discovering unlabeled positives (**Fig. 3a, 4a**). We ascribe this observation to the effect of GNN layers to regularize the ML training by weighting adjacent node's features and acting as a low pass filter, and thus lowering the risk of overfitting to known positive examples. However, the performance of different GNN layers varies greatly, and in fact several decrease overall model performance (**Extended Data Fig. 5**), whereas others achieve high prediction performance also for novel core genes (**Fig. 3d**). N2V+MLP works well for known positives but underperforms for unknowns (**Fig. 3c**). Since we are predominantly interested in the discovery of unlabeled positives, GNNs are the more promising choice. However, GCN[46] and RGCN[48] layers generally underperform (**Fig. 1b**). On the one hand GCN, which mainly exploits the features of all neighboring nodes, is outperformed by an MLP, and on the other hand N2V+MLP, which exploits edges but no neighboring node features outperforms the MLP. So, It appears that the node features of neighbors are not helpful, while the edges are *per se* informative. This explains that TAG[47] ameliorates the dip in performance caused by the GCN layer by incorporating skip-connections, which can bypass the message passing. In this setting, the TAG layer can be understood as a conditional GNN layer which, if the adjacent node's features are unhelpful, can revert to an MLP for any given node. This is also mirrored in the fact that TAG always performs at least as good as an MLP if the network topology is unhelpful (**Fig. 1b**)**.** However, this mechanism neglects the information contained in the topology of the graph alone, i.e. in the edges themselves, which appear to be helpful as indicated by the N2V+MLP performance. Conceptually, the FiLM[49] layer, does not only use the sender node's features for its convolution, but also the receiving nodes' features and the type of the connecting edge. When we examined which features are most important for FiLM predictions we found that the influence of the adjacent node's input features was negligible for individual examined nodes (see **Supplementary Fig. SF1),** and globally the latent features of surrounding nodes make up only a tiny proportion of all messages in the FiLM layer (**Supplementary Fig. SF2, Supplementary Note 5)**. This indicates that also for FiLM the receiving node's features and the edge type are the most important factors determining the message (**Supplementary Note 6**). Thus, reminiscent of the N2V+MLP results, the FiLM layer predominantly learns the topology free of the influence of adjacent node's features but has the additional advantage of incorporating edge types.

The fact that FiLM's messages are mostly dependent on the receiving node's features and the edge type is also reflected in the results shown in **Fig. 5**. The importance of the edges is almost identical between edges of the same type, with only minute differences caused by the sending node's features. It also indicates that only a handful of roughly 300 incident edges is important for the prediction. Collectively these analyses indicate that methods gain performance when they have the option to ignore adjacent nodes' features and instead learn patterns of select incident edges. Biologically, it is possible that neighborhood functions are encoded in the network topology, e.g. in the form of protein complexes, or a sufficiently specific interaction wiring in different modules, and thus learned indirectly by the GNNs as a pattern that better captures functions than individual node features. In addition, the wealth of connections, and network incompleteness, likely also impact on the observed phenomena. Novel methods that are designed to distill the information of edges[138,139] or topological features[140,141] of the graph alongside the node features could therefore be a valuable addition to future iterations of comparable work.

**Supplementary Note 4: Impact of network biases on predictions**

As the bias of aggregating small-scale literature is visible in network characteristics concerning Mendelian disorder genes (**Supplementary Note 1**),we monitored how these biased inputs affect the predicted candidate genes and potentially inhibit new insights. TAG, which is trained only on IntAct Direct Interaction, is heavily biased towards predicting genes that have a degree larger than zero in the IntAct Direct Interaction network (**Extended Data Fig. 8a**) and in the case of immune dysregulation even exclusively predict such genes as candidates (**Extended Data Fig. 8b**). This indicates that presence in the IntAct direct network, and hence the fact that the involved proteins were deemed 'interesting' by researchers to justify their biochemical purification and *in vitro* interaction studies, and hence previous perceptions of a gene's importance, was a key feature in their prediction by TAG. The argument that the scientific communities' accumulated knowledge reflected in such a focused deeper characterization of relatively few genes corresponds to underlying biological importance has previously been refuted[18,25,26]. The candidate genes predicted by FiLM, which is trained on all networks simultaneously including IntAct Direct Interaction and is aware of edge types, shows much lower odds ratios (OR) for genes from IntAct Direct Interaction, but still significantly higher than 1 (**Extended Data Fig. 8a)**. The Node2Vec in N2V+MLP is also trained on all networks simultaneously but is not aware of edge types, so IntAct Direct Interaction makes up only 0.36% of all edges, thus further reducing the ORs of genes contained in IntAct Direct Interaction among its candidates. Importantly, when FiLM is trained on all networks except IntAct Direct Interaction and IntAct Physical Association (FiLM Unbiased), we still reap the benefits of GNNs without biasing predictions towards genes included in IntAct. However, even the candidates produced by N2V+MLP and FiLM Unbiased are weakly enriched for genes involved in IntAct Direct Interaction, reflecting a bias of IntAct Direct Interaction for immune system regulation. This interpretation is supported by the validation of FiLM Unbiased predictions using mouse KOs, in which the OR drops for immune dysregulation, compared to the initial FiLM predictions, but not for other disease groups like cardiovascular disease (**Extended Data Fig. 9a, Supplementary Table ST11)**. Furthermore, differentially expressed genes and drug targets (**Extended Data Fig. 9b**, **c**) show comparable levels of enrichment for most diseases whether IntAct Direct Interaction is used with FiLM or not. Intriguingly, for immune dysregulation the enrichment of differentially expressed genes increases when IntAct data are removed, indicating that these networks might also be biased towards genes examined in small-scale mouse experiments, which is consistent with the reasoning that laborious mouse knock-out and in vitro studies are more readily done for genes/proteins considered important. Moreover, for immune dysregulation only candidates produced by the unbiased version of FiLM are enriched for druggable genes which are not yet drug targets (**Extended Data Fig. 9c,** Dr-), indicating that the biases inherited from small scale literature could indeed prevent the discovery of new drug development opportunities.

**Supplementary Note 5: Importance of neighborhood node features**

We have shown that some edges in the direct neighborhood of nodes and a selection of input features are vital for classification (**Fig. 5**). Besides their own input features, other node's input features appear to be irrelevant for the prediction (**Supplementary Fig. SF1**). Some of the genes (e.g. HBB) are not even in the 2-hop neighborhood of the query node, indicating that the threshold of 0.1 (dotted gray line) is a good threshold to separate signal from statistical noise. The fact that TNFRSF25 and TNFRSF6B's input features do not seem relevant, even though their connection to TNFSF15 is relevant (**Fig. 5**) indicates that it is in fact the physical connection of the proteins (i.e. the edge), and not the neighboring node's features, that is meaningful. It furthermore underlines the power of the FiLM layer, which can override the message coming from these nodes and thus can create a meaningful message from unhelpful incoming node features (see Methods Section for details). Since the overridden message is conditioned only on the receiving node and the connecting edge type, the message is identical for all incoming connections of the same type. This explains why the edge importance of edges of the same type are almost identical (**Fig. 5**) and why RGCN, which lacks this mechanism and relies on transformations of the sender's features, fails to perform adequately compared to FiLM (**Fig. 1b**).

To further investigate the conjecture that neighboring node's features are mostly irrelevant for the prediction while the edges themselves are relevant, we have ascertained the influence of sender, receiver and edge on the message passing dynamic of the FiLM layer. The FiLM layer introduces an offset beta and a linear coefficient gamma for every feature of an incoming message $x_u^{(t)}$ from the sender node $u$ in the neighborhood of $v$ based on the edge type $r$ and the receiver node $v$:

$$x_v^{(t+1)} = \sum_{r \in R} \sum_{u \in N(v)} \sigma(\gamma_{r,v}^{(t)} \odot W_r x_u^{(t)} + \beta_{r,v}^{(t)})$$

Thus, the influence of the neighborhood node's features is only relevant for the first part of the term:

$$\gamma_{r,v}^{(t)} \odot W_r x_u^{(t)} ,$$

while the bias $\beta_{r,v}^{(t)}$ is only dependent on the receiver node $v$ and the edge type $r$. We can therefore assess the balance between the influence of the neighborhood node's features and the features of the receiving node the following ratio:

$$\frac{\gamma_{r,v}^{(t)} \odot W_r x_u^{(t)}}{\beta_{r,v}^{(t)}} ,$$

which is close to 0 if the message is dominated by the bias term $\beta_{r,v}^{(t)}$ and thus irrelevant of the neighborhood node's features. A high value still does not guarantee a high influence of $x_u^{(t)}$, but we can assume that it is increasingly relevant if the model decides to modulate it via $\gamma_{r,v}^{(t)}$ instead of simply overriding it via $\beta_{r,v}^{(t)}$.

The message features passed along the edges of all networks of the first FiLM-layer are, in fact, dominated by the bias term $\beta_{r,v}^{(t)}$, rendering the sender's latent features irrelevant (**Supplementary Fig. SF2a)**. The message features passed along the edges of gene regulatory layers by the second layer (see **Supplementary Fig. SF2b**) are less heavily dominated by the bias term. However, this still does not mean that the actual input features are important, since the latent features of the senders have already been influenced by their incident edges in the first layer. It does, however, indicate that protein-protein networks and gene regulatory networks convey different notions of neighborhoods, which might be influenced by the former being bidirectional and the latter being unidirectional.

**Supplementary Note 6: Additional predicted examples**

Analogous to the immune dysregulation examples in the main text, we also explored candidate genes predicted by FiLM for cardiovascular disease for suitable targets for drug development. OBSCN and ITGA7 receive high Consensus Scores (11 and 9, respectively) and their protein products are druggable but not yet targeted by any drug.

OBSCN encodes the protein obscurin is a large, modular protein with more than 80 exons and 28 transcript isoforms[142], which fulfill a wide range of functions in different tissues including skeletal and heart muscle[143]. Specific mutations in the OBSCN gene are implicated in hypertrophic cardiomyopathy[144], age-dependent cardiac remodeling and arrythmia[145]. Furthermore, obscurin has been implicated in non-muscular functions and pathologies[143]. Film bases its prediction of OBSCN as candidate gene on its location downstream of multiple transcription factors across several tissues, and implicating it in inflammation (STAT2, ZNF384)[146–148], angiogenesis (BRF1)[127], and immune dysregulation after ischemic damage (IRF3)[150] (**Extended Data Fig. 10a)**.

The integrin subunit alpha 7 encoded by ITGA7 is located in the cell membrane and involved in cell-cell and cell-matrix communication, and has been implicated in migration and invasion of malignant cells in metastasis formation[151,152]. Recently it was shown that mutations in ITGA7 contribute to congenital muscular dystrophy[153], adult-onset cardiac dysfunction[154] and cardiomyopathy[155,156], implicating a role in the etiology of cardiovascular diseases. The fact that FiLM bases its prediction on ITGA7 being located downstream of the estrogen related receptor alpha, encoded by ESRRA (**Extended Data Fig. 10b)**, opens the possibility that this gene mediates sex-related differences in genetic cardiomyopathies[157].

Neither OBSCN nor ITGA7 have been detected in GWAS studies for any heart-related traits, except PR interval (GCST010321) in the case of OBSCN. Despite this lack of detection, some OBSCN variants are known to contribute to left ventricular compaction[158] and dilated cardiomyopathy[159]. Despite the absence of GWAS signal, Speos identified them as core gene candidates based on their tissue-specific gene expression (**Extended Data Fig. 10c, d)**. Especially their high expression in the left ventricle and atrial appendage are vital for their classification as expected for factors contributing to cardiovascular disease, as these anatomical regions are key players in several related pathophysiologies[158,160]. Although the understanding of the role the two genes play in cardiovascular disease is still in its infancy, at least OBSCN already raised expectations for novel treatments and therapeutics[143], underscoring the value of Speos' predictions for hypothesis development, even when significant genome wide associations have not been detected.

### Supplementary Figures

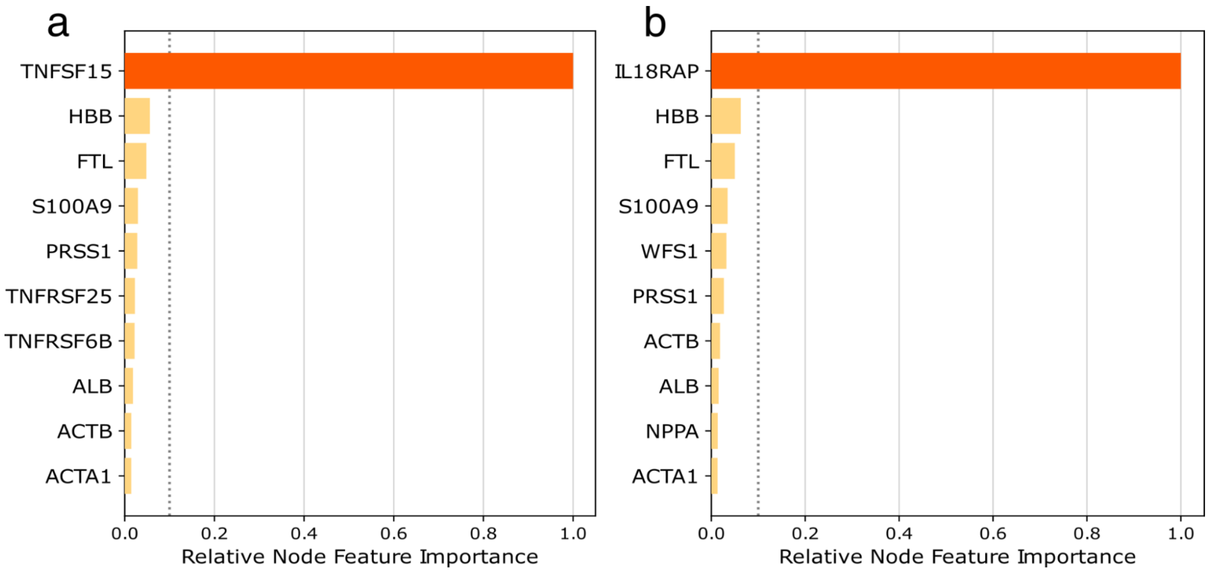


**Supplementary Fig. SF1 | Neighborhood Feature Importance**. Shown is the cumulative normalized importance of a node's input features for the prediction of query nodes **a**: TNFSF15 and **b**: IL18RAP as candidate genes for immune dysregulation predicted by FiLM. Shown are the top 10 most influential nodes. A node that has the most relevant input features for the prediction of the query nodes every for model in the ensemble has a value of 1, while a node that has the least relevant input features for every model has a value of 0. The dashed gray line denotes the threshold of 0.1.

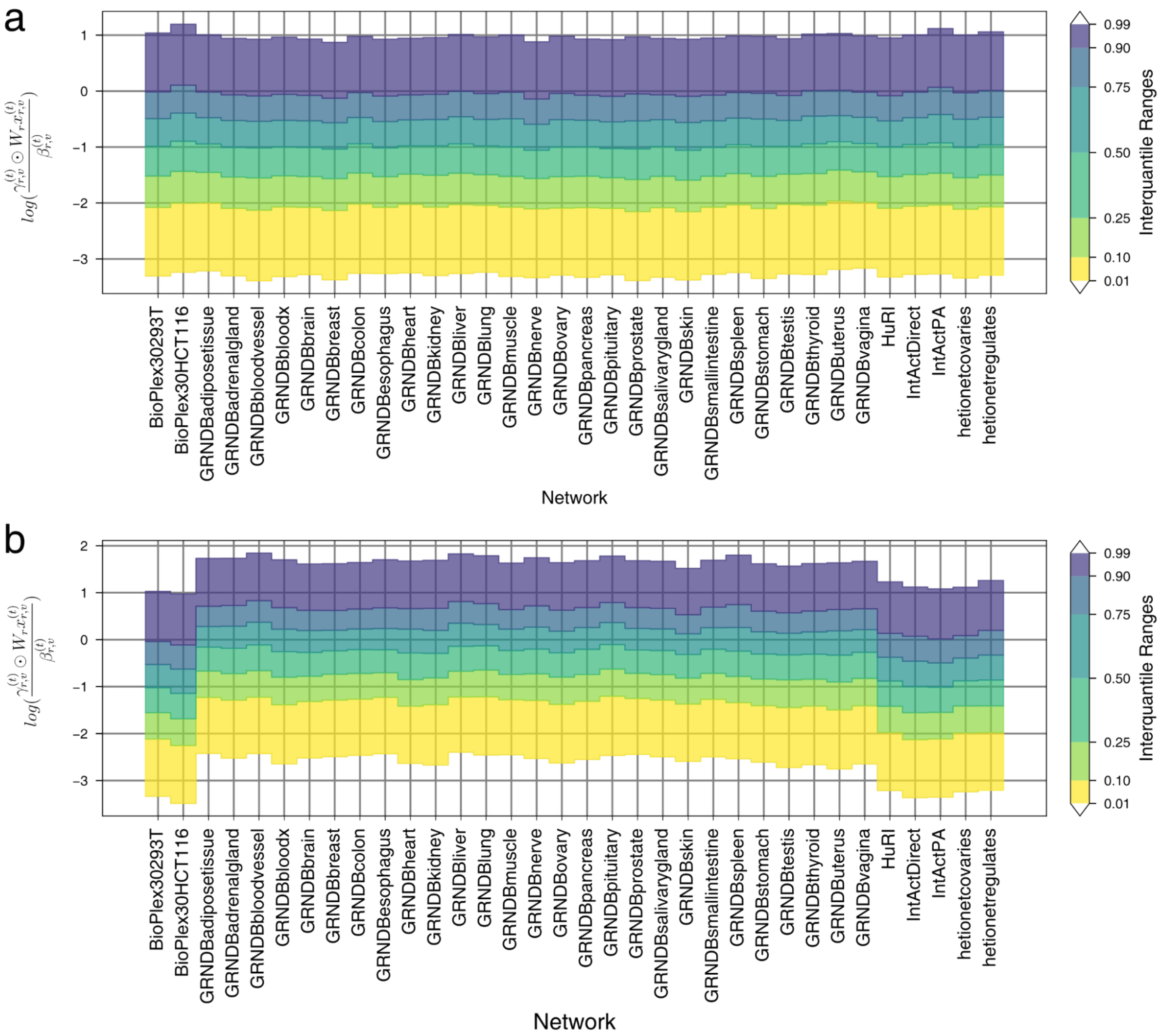


**Supplementary Fig. SF2 | Ratio of Sender to Receiver features.** Shown is the influence of the bias term on the features of the messages passed along the edges of all networks in a single FiLM model trained for immune dysregulation. **a,** values for the first layer. **b** shows the values for the second layer. Low values indicate that the message is dominated by the bias term and thus independent of the sender's features.
